# Rare Variant Phasing and Haplotypic Expression from RNA-Sequencing with phASER

**DOI:** 10.1101/039529

**Authors:** Stephane E. Castel, Pejman Mohammadi, Wendy K. Chung, Yufeng Shen, Tuuli Lappalainen

## Abstract

Haplotype phasing of genetic variants is important for clinical interpretation of the genome, population genetic analysis, and functional genomic analysis of allelic activity. Here we present phASER, a fast and accurate approach for phasing variants that are overlapped by sequencing reads, including those from RNA-sequencing (RNA-seq), which often span multiple exons due to splicing. This provides 1) dramatically more accurate phasing of rare and *de novo* variants compared to population-based phasing; 2) phasing of variants in the same gene up to hundreds of kilobases away which cannot be obtained from DNA-sequencing reads; 3) high confidence measures of haplotypic expression, greatly improving power for allelic expression studies.

---

The phasing of rare and **de novo** variants is crucial for identifying putative causal variants in clinical genetics applications, for example by distinguishing compound heterozygotes from two variants on the same allele. Existing methods to phase variants include phasing by transmission ^1^, only available in familial studies, population based phasing ^2,3^, which is ineffective for rare and *de novo* variants, phasing by sequencing long genomic fragments ^4,5^ which requires specialized and costly technology, and phasing using expression data by inferring haplotype through allelic imbalance ^6^, which only applies to loci with well-detected allelic expression. An alternative approach termed “read backed phasing” uses readily available short read DNA-sequencing (DNA-seq) ^8–10^, however it is limited by the relatively short distances which can be spanned by the reads. Our approach, called phASER (<u>ph</u>asing and <u>a</u>llele <u>s</u>pecific <u>e</u>xpression from <u>R</u>NA-seq), extends the idea of read backed phasing to RNA-seq reads, which due to splicing enables phasing of variants over long genomic distances. Data from both DNA-seq and RNA-seq libraries can be integrated by phASER to produce high confidence phasing of proximal variants, primarily within the same gene, and when available population phasing can also be leveraged to produce full chromosome-length haplotypes (Figure 1a). Source code and complete documentation for phASER and its associated tools are available through github (https://github.com/secastel/phaser).

Assembling haplotypes from observations of alleles on the same read is a necessary step of read backed phasing, and has been accomplished using various approaches ^8–10^. Our approach in phASER employs a two step method, first defining edges between the alleles of each pair of variants observed on the same sequencing fragments, and second, determining the most likely phase within a set of connected variants given the edges defined in the first step (Figure S1). During the first step the phase with the most supporting reads is chosen, and a binomial test is performed to determine if the number of reads supporting alternative phases is greater than would be expected from sequencing noise, allowing for filtering of low confidence phasing (Figure S1a, Figure S2a). For the second step, phASER counts the number of edges that support each possible haplotype configuration (2^n^ variants), and selects the configuration with the most support. To prevent an exponential increase in haplotype search space while maintaining accuracy phASER splits large haplotypes into sub-blocks at points spanned by the fewest edges (Figure S1c, Figure S2c). Phasing is performed chromosome wide, with no restriction on the distance between variants, which allows phasing at the longer genomic distances spanned by RNA-seq reads.

As a gold standard we compared phASER used with high coverage RNA-seq data generated from a lymphoblastoid cell line (LCL) ^11^ to Illumina’s NA12878 Platinum Genome, sequenced at 200x and phased by transmission using parental data (http://www.illumina.com/platinumgenomes/), and found that with default settings phASER identified the correct phasing for 98% of the variants phased. Unlike population based phasing, read backed phasing across sequencing assays performed well for low frequency variants (Figure 1b). Finally, we benchmarked haplotype assembly in phASER against HapCUT ^9^ and the GATK Read Backed Phasing tool, which work only for DNA-seq reads, using both simulated and experimental WGS data as well as WES data. We found phASER to be similar in accuracy, runtime and haplotype length to HapCUT (Figure S3), while having additional features required for RNA-seq based phasing. Both phASER and HapCUT were dramatically more accurate than the GATK tool (Figure S3a).

In order to evaluate the increase in phasing distance facilitated by RNA-seq reads, we compared phASER results between WES, WGS, and two read lengths of paired-end (PE) RNA-seq (75bp and 250bp) from 4 Genotype Tissue Expression (GTEx) individuals where matched libraries were available ^12^. As expected, long read RNA-seq yielded the greatest proportion of distantly phased variants, with an average of 4300 equal to 5.8% of variants phased greater than 5kb, while at this distance WES and WGS phased 0 and 7 variants respectively (Figure 1c). At large distances performance of RNA-seq phasing decreased as a result of read mapping errors, however this could be easily addressed by filtering reads based on alignment score (Figure S4a). Using RNA-seq reads phasing remained accurate over a range of read lengths (Figure S4b), but longer read lengths greatly increased both the distance and number of variants that could be phased (Figure S4c).

When population phased data are available, haplotype blocks are phased relative to each other, producing a single genome wide phase through a method we call phase anchoring. Phase anchoring uses the population phase of each variant in a block weighted by their allele frequencies to assign a genome wide phase, since common variants are more likely to have correct population phasing (Figure S2b). A similar approach is used by methods that integrate read backed phasing with population phasing ^10^, however including it in our method allows this strategy to be used with RNA-seq reads and prevents the need to perform population phasing each time new sequencing data for a sample is available. Using our approach with RNA-seq from accessible tissues enabled genome wide phasing of up to 15.4% of rare coding variants (MAF ≤ 1% in GTEx), and 21.3% when tissues were combined, while WES yielded 19% and 5× WGS yielded 11.1% (Figure 1d). When used in combination, the addition of combined RNA-seq data enabled a 1.5× increase in phasing for WES and a 2.4× increase in phasing for WGS. When considering all rare variants, WGS performed better, and the contribution of RNA-seq to WES was more significant (Figure 1e).

We next sought to benchmark phASER when used with RNA-seq data in the context of genetic studies. First we used GTEx data to demonstrate the number of coding variants that could be phased as a function of the number of tissues for which RNA-sequencing data is available. We began with whole blood, and progressively added libraries from up to 14 other distinct tissues. With joint phasing using 14 tissues, almost 50% of all heterozygous coding variants could be phased with at least one other variant (Figure S5a). When used individually, the total proportion of coding variants that could be phased for a given tissue was 15–27% (Figure S5b), and was dependent on transcriptome diversity ^13^ (Figure S5c), but not total read depth (Figure S5d).

We next applied phASER to assess its ability to identify cases of compound heterozygosity for damaging variants using exome and RNA-seq data from 345 1000 Genomes individuals ^14,15^. First we assessed the accuracy of compound heterozygote calls using population phasing compared to exome + RNA read backed phasing. As expected, protein-truncating and splice variants that are usually rare were enriched in cases where population phasing was incorrect, with cases involving stop gain variants being 2.9× more likely to be phased incorrectly than others (Figure S6). Next, to demonstrate the advantage of using RNA-seq data over exome alone for phasing, we identified instances of compound heterozygosity involving at least one rare (MAF < 1%) variant with predicted loss of function (LoF) or damaging protein effects ^16^. Including RNA-seq data from only one tissue (LCLs) ^14^ increased the number of compound heterozygotes that could be identified in the most severe class (LoF × Damaging) by 1.3× over WES alone, and ranged between 1.19 – 1.15× for other combinations, demonstrating the added benefit of phasing over larger distances (Figure 2a).

Finally, we used paired exome and fibroblast RNA-seq from 20 patients with congenital diaphragmatic hernia ^17^ to illustrate phASER’s application to a typical clinical workflow to prioritize putatively causal variants and benchmark the advantage of including RNA-seq reads for phasing. Assuming that causative variants would be rare, recessive, damaging, and expressed in the tissue of disease relevance, phase information generated with phASER from exome and RNA-seq reads could prioritize a median of 25 alleles involved in cases of compound heterozygosity per individual (*trans*), while assigning a lower priority to a median of 44 alleles, where the alleles where on the same haplotype (*cis*) (Figure 2b). The inclusion of RNA-seq boosted the number of cases that could be identified by 2.6× for *trans* and 1.4× for *cis* interactions.

Outside of medical genetics, variant phasing is important for allelic expression (AE) analysis, which aims to quantify the relative expression of one allele versus another, and has emerged as a powerful method to study diverse biological processes including cis-regulatory variation ^14^, parent of origin imprinting ^18^, and protein-truncating variants ^19^. AE is typically measured at single heterozygous variants, however the unit of interest is often a gene or transcript, which may contain many variants. Integrating read counts across phased variants can greatly improve the power to detect AE, but simply adding up allele counts can lead to double counting of reads (if variants are covered by the same read), and both false positives and negatives as a result of incorrect phasing. To address this limitation, when used with RNA-seq data, phASER quantifies and reports the expression of phased haplotypes by reporting the number of unique reads that map to each. To benchmark the impact of this utility we generated haplotypic counts at genes with known expression quantitative trait loci (eQTL) ^14^ for 345 Geuvadis samples using either single variants with population based phasing alone, or phased haplotype blocks generated by phASER (Figure S7). By improving phase and preventing double counting phASER reduced false positives at 56.2% of genes tested, while uncovering false negatives at 7.3% (Figure 2c).

In summary, phASER provides scalable and high confidence variant phasing, incorporating RNA-seq and DNA-seq data with population phasing, allowing phasing over longer distances than previous read based methods. We have demonstrated that this method has direct applications in medical genetics, where improved resolution of compound heterozygotes can lead to changes in their interpretation. Furthermore, phASER improves the accuracy of haplotypic expression when integrating allelic counts across variants by reducing false positives. Our approach will complement the existing repertoire of phasing methods ^3^ and makes use of a readily available experimental data type that has become trivial to produce, allowing for phasing of rare and distant variants at high accuracy. As RNA-seq experiments become commonplace in medical and population scale studies, phASER will become a valuable tool for rare variant phasing.

**Figure 1.**
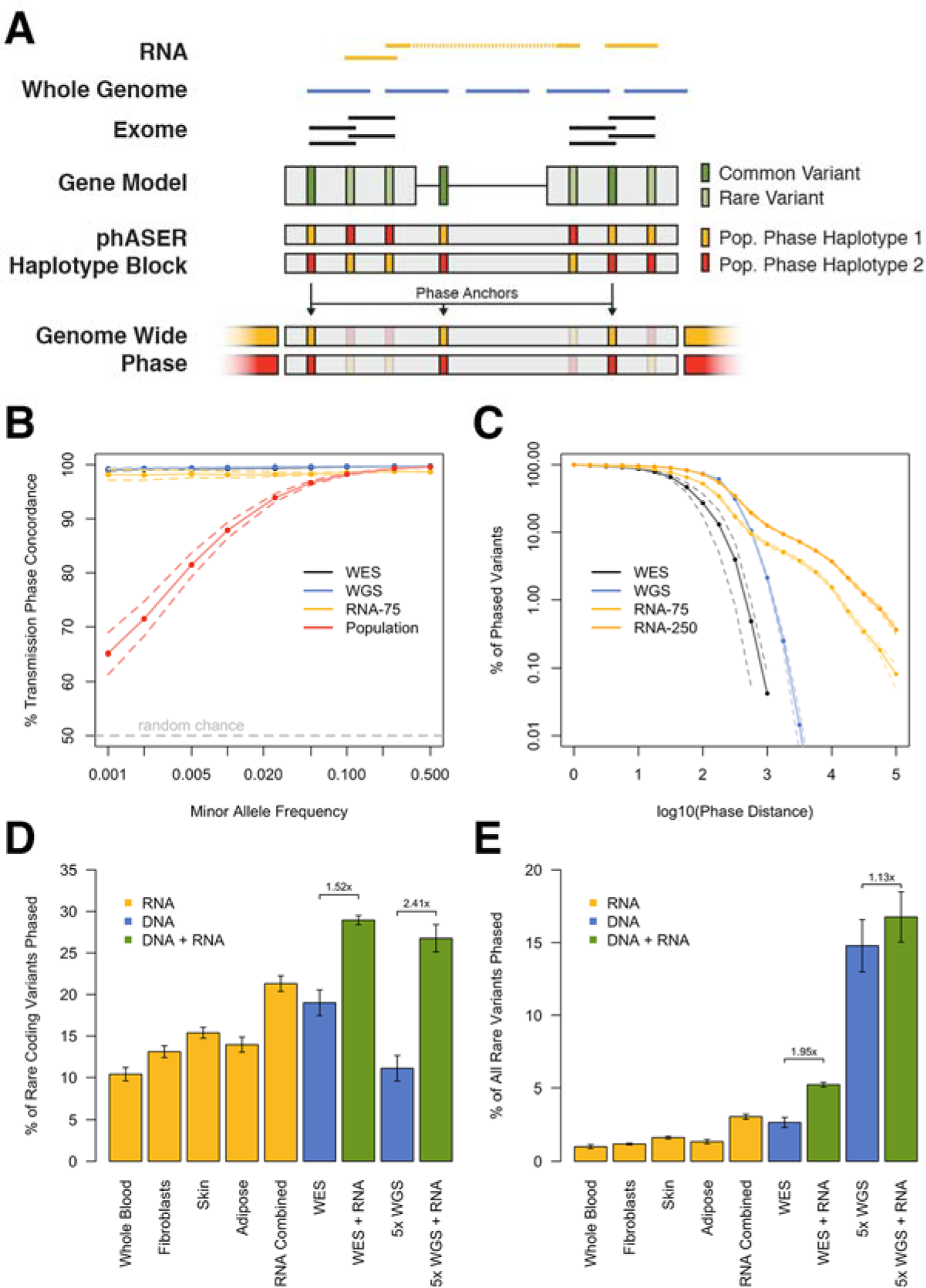
Read backed haplotype phasing that incorporates RNA-seq using phASER. A) phASER produces accurate variant phasing through the use of combined DNA and RNA read backed phasing integrated with population phasing. Due to splicing, RNA-seq reads often span exons and UTRs, allowing read backed phasing over long ranges, while high coverage exome and whole genome sequencing can phase close proximity variants. A local haplotype is produced by testing all possible phase configurations, and selecting the configuration with the most support (Figure S1). Local haplotype blocks can be phased relative to one another when population data is available by anchoring the phase to common variants, where the population phase is likely correct. B) Concordance of read backed phasing across sequencing assays and population phasing with phasing by transmission using the Illumina NA12878 Platinum Genome as a function of variant minor allele frequency. Concordance is defined per variant as the percentage of variant - variant phase events that are correct as compared to the known transmission phase. C) Percentage of phased variants that can be phased at greater than or equal to increasing genomic distances using WES, WGS, paired-end 75 and 250 RNA-seq data in two tissues (whole blood and LCLs) of four GTEx individuals. Solid lines represent the means, and dotted lines the standard error. D-E) Contribution of read backed phasing at rare coding (MAF ≤ 1%) variants (D) and all rare variants (E) across sequencing assays and GTE× RNA-seq tissue types for four individuals. Values shown are the mean percentage of rare variants within an individual that can be assigned a genome wide phase using phase anchoring. Error bars show the standard error. The fold increase in the number of rare variants that can be phased using DNA-seq with the addition of combined RNA-seq libraries is indicated.

**Figure 2.**
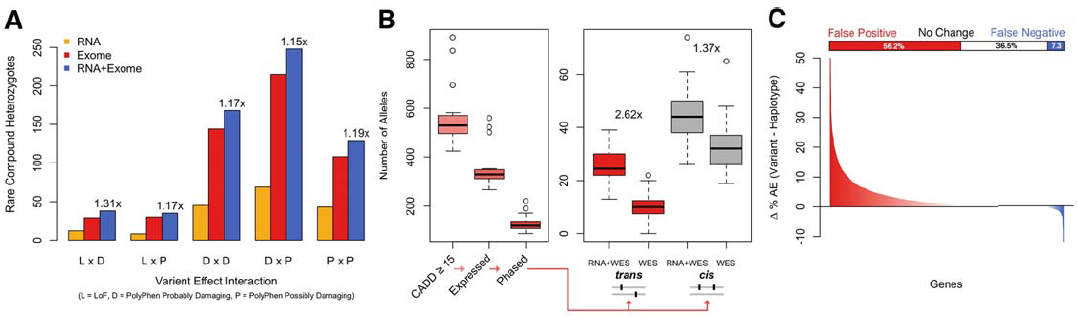
Application of RNA-seq based haplotype phasing to studies of functional variants and allelic expression analysis. A) Instances of compound heterozygosity involving rare (MAF < 0.01) loss of function (L), probably damaging (D) or possibly damaging (P) coding variants called using phase data generated by phASER with either RNA-seq reads, exome-seq reads, or both for 345 1000 Genomes European individuals with Geuvadis LCL RNA-seq data. The fold increases in the number of compound heterozygotes when RNA- seq data is included are indicated. B) Example application of phASER to prioritize rare (alternative AF < 0.01 in 1000 Genomes) recessive alleles in a clinical study that includes both exome-seq and RNA-seq in a tissue of clinical relevance ^17^ Boxplots show the number of heterozygous alleles per individual after these successive filtering steps were applied: CADD phred score ≥ 15, expressed in fibroblast RNA-seq data, phased with read backed phasing, involved in either *trans* or *cis* interactions with another deleterious variant (CADD ≥ 15) using RNA and exome data (RNA+WES) or exome alone (WES). The fold increases from including RNA-seq data are indicated. C) The difference in percentage of individuals with significant (binomial test, q < 0.05) allelic expression for each gene with a known heterozygous *cis* eQTL calculated by either summing all single variant read counts across haplotypes using population phasing, or by summing phASER haplotype blocks phased relative to each other with phase anchoring (see Figure S7). Genes where an increase in the percentage of individuals with significant allelic expression is observed are colored red, representing false negatives when summing single variant counts, while those with a decrease, representing false positives, are colored blue. The bar plot above indicates the percentage of the 1118 genes where allelic expression was measured that fall into each category.

## Methods

### Implementation of read backed phasing in phASER

phASER is written in Python and requires the following libraries: IntervalTree, pyVCF, SciPy, NumPy. In addition it requires Samtools and Bedtools to be installed. Aligned reads (BAM format) are mapped to heterozygous variants, and for each heterozygous variant a hashed set of all overlapping reads is produced which allows for quick comparison of overlapping reads between variants. Connections between variants are established whenever a read (or read mate) overlaps more than one variant. For each pair of connected variants edges between alleles are defined by determining the phase that has the most read support, and a test i performed to determine if there is significant evidence of a conflicting phase (see below) (Figure S1a). Those edges that fail, based on a user defined significance threshold (by default nominal p < 0.01) are removed. (It should be noted that this test addresses instances where due to sequencing error individual bases on a read have been misread, and it does not address errors with read mapping, which can happen at sites with genetic variation. For a discussion of methods to address allelic read mapping issues please see Castel et al., 2015) Haplotype blocks are generated by starting with a single unphased variant and recursively adding all othei variants with read connections. Block construction is completed when no further variants can be added. Once all haplotype blocks have been generated, phasing of the variants is determined using the previously defined allele edges. If the allele edges within a haplotype block resolve into two distinct groups, where each group contains only one allele of a given variant, the haplotype is considered to be conflict free and the phase is reportec (Figure S1b). In instances where two haplotypes cannot be immediately resolved, the haplotypic configuration with the most edges supporting it is identified. This is accomplished by testing all possible haplotype configurations (2^n^), however runtime is prevented from exponentially increasing by splitting haplotypes intc sub-blocks at positions spanned by the fewest number of edges (Figure S1c–d). These sub-blocks are then phased relative to one another to produce a single haplotype phase (Figure S1e). The maximum number of variants within a sub-block is user defined, and is 15 by default. Our simulations show that when using this approach accuracy remains high, while runtime is drastically reduced (Figure S2c). Read variant mapping, edge definition and haplotype assembly can all by parallelized for an increase in speed (Figure S3b).

### Statistical test for conflicting phasing between two variants

For each SNP pair covered by at least one read a test is performed that determines if the number of reads supporting alternative phasing (any phase other than the configuration chosen by phASER) could be observed by chance from sequencing noise alone. In this test, significance indicates more conflicting reads than would be expected from noise alone, and thus suggests that there may be an error in the phase selected by phASER. A conservative approach is to filter out any variant connections with a nominal p value of < 0.01 (see Figure S1b) Lower p value thresholds can be used to retain more blocks, with the caveat that some may have incorrect phasing. Filtered connections will not be used during the haplotype block construction process.

The test is based on a uniform error model in which a true allele nucleotide can be substituted randomly to any other nucleotide. All pairwise substitutions in this model are assumed to be equally probable. We denote this pairwise substitution probability with ε. Let us assume a pair of SNPs a_1_|a_2_ and b_1_|b_2_ with a haplotype structure a_1_b_1_| a_2_b_2_. Let a_1_:b_1_ denote a read spanning alleles a_1_ and b_1_. Reads supporting the true haplotype in this case are a_1_:b_1_, and a_2_:b_2_, and any other configurations correspond to conflicting evidence. Let us only consider reads generated from the first haplotype. The probability of observing a read supporting the correct haplotype structures, *p_s_*, is (1 − 3*ε*)^2^ + *ε*^2^, where the first term corresponds to the probability of observing a_1_:b_1_ (the case in which either of the sites being affected by noise in the read), and the second term is the probability of observing a_2_:b_2_ (the case in which both sites were altered by noise in the read and happened to have the other second allele). Binomial distribution is used to evaluate the probability of observing equal or less supporting reads for two given SNP sites:

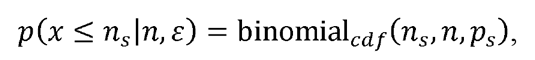

where *n_s_* and *n* are the number of reads supporting the chosen haplotype structure and total number of reads respectively. The pair-wise substitution rate *ε* is calculated from over all SNP sites as:

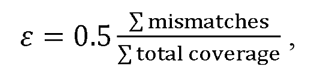

where mismatches correspond to all cases that a nucleotide other than the reference or the alternative allele where observed at the site. Only variants where > 50% of the reads come from the reference and alternative alleles are used to calculate the substitution rate to avoid inflation of noise estimates as a result of genotyping error.

### Genome wide phasing using phase anchoring

When population phased data is available phASER will attempt to determine the genome wide phase for each haplotype block using phase anchoring. Phase anchoring operates on the assumption that common variants are more likely to be phased correctly using population data, so for each variant within a haplotype the genome wide phase determined by population phasing is weighted by the variant allele frequency. The genome wide phase across all variants with the most support after weighting is selected for each haplotype block.

For each haplotype block:

*α* (phase support for configuration 1) = Σ(MAF of variants supporting configuration 1)
β (phase support for configuration 2) = Σ(MAF of variants supporting configuration 2)
if α > β genome wide phase = configuration 1
if α < β genome wide phase = configuration 2
Anchor Phase Confidence = max(α, β) / (α+β)

### phASER settings

The following settings were used unless otherwise noted. For all libraries: alignment score quantile cutoff of 0. 05, BASEQ of 10, and conflicting configuration threshold of 0.01, indels ignored, and maximum block size of 15. For RNA-seq libraries: no maximum insert size, MAPQ of 255 (indicates unique in STAR). For exome-seq libraries: 500 bp maximum insert size, MAPQ of 40. For whole genome sequencing libraries: 1000 bp maximum insert size, MAPQ of 40. All variants in HLA genes were filtered.

### Code availability

Source code and complete documentation for phASER and its associated tools are available through github (https://github.com/secastel/phaser). In addition to the phASER core software we provide two scripts: one which given an input VCF that has been phased using phASER will identify all interactions between alleles (phASER Annotate), and retrieve information such as allele frequency and predicted variant effect if supplied with the appropriate files, and second, a script which will use haplotypic counts produced by phASER in combination with population phasing to produce gene level haplotypic read counts for use in allelic expression studies (phASER Gene AE).

### Data processing, usage and availability

For analyses involving 1000 Genomes individuals, phase 3 genotypes and population phasing where used with hg19 aligned exome-seq data, both downloaded from the 1000 Genomes website (http://www.1000genomes.org). Raw (FASTQ) RNA-sequencing data from 1000 Genomes individual derived LCLs was downloaded from the European Nucleotide Archive (ERP001942), and aligned with STAR to hg19. For comparison of phase statistics between sequencing experiments the following GTEx individuals were used: S32W, T5JC, T6MN, WFON. Both short and long read RNA-seq was obtained for whole blood and LCLs, and aligned using STAR to hg19. WES reads were aligned using Bowtie 2 to hg19. GTEx data is available through dbGaP for authorized users (phs000424.v6.p1). For rare variant phasing comparison, RNA-seq from whole blood, fibroblasts, sun exposed skin, and adipose were used, alongside WES and WGS libraries, from GTEx individuals X4EO, XUW1, U8XE, XOTO. GTEx individual ZAB4 was used for comparison of number phased variants versus number of RNA-seq tissues used. For comparison to transmission phasing the following data from the 1000 Genomes individual NA12878 was used: exome-seq downloaded from 1000 Genomes website, whole genome sequencing data (NCBI SRA ERS179577), RNA-seq from a LCL (NCBI GEO GSM1372331), transmission phased genotypes (Illumina Platinum Genome, http://www.illumina.com/platinumgenomes/). Whole genome sequencing libraries were down sampled to 5× to increase speed of analyses and ensure comparable read depths across sequencing assays.

### Benchmarking

Benchmarking was run on CentOS 6.5 with Java version 1.6 and Python 2.7 on an Intel Xeon CPU E7- 8830 © 2.13 GHz, with GATK v3.4, HapCUT v0.7, and phASER v0.5. The GATK tool was run with default settings, with the exception of: min_mapping_quality_score = 40, maxPhaseSites = 15, min_base_quality_score = 10. HapCUT was run with the following settings: maxIS = 500 (WES) or 1000 (WGS), mbq = 10, mmq = 40. phASER was run with default settings (see phASER settings). Simulated PE 75 WGS data was produced with ART Chocolate Cherry Cake 03–19–2015 ^20^ from a NA12878 1000 Genomes Phase 3 population phased reference. WES and WGS libraries used were those listed above.

## Acknowledgements

We would like to thank the Geuvadis Consortium, the GTEx Consortium, the members of the Lappalainen lab, and the bioinformatics team of the New York Genome Center. The Genotype-Tissue Expression (GTEx) Project was supported by the Common Fund of the Office of the Director of the National Institutes of Health (commonfund.nih.gov/GTEx). Additional funds were provided by the NCI, NHGRI, NHLBI, NIDA, NIMH, and NINDS. Donors were enrolled at Biospecimen Source Sites funded by NCI\SAIC-Frederick, Inc. (SAIC-F) subcontracts to the National Disease Research Interchange (10XS170), Roswell Park Cancer Institute (10XS171), and Science Care, Inc. (X10S172). The Laboratory, Data Analysis, and Coordinating Center (LDACC) was funded through a contract (HHSN268201000029C) to The Broad Institute, Inc. Biorepository operations were funded through an SAIC-F subcontract to Van Andel Institute (10ST1035). Additional data repository and project management were provided by SAIC-F (HHSN261200800001E). The Brain Bank was supported by a supplement to University of Miami grant DA006227. Statistical Methods development grants were made to the University of Geneva (MH090941), the University of Chicago (MH090951 & MH090937), the University of North Carolina - Chapel Hill (MH090936) and to Harvard University (MH090948).

We are thankful to all the CDH families for their generous contributions. We are grateful for the technical assistance provided by Patricia Lanzano, Jiancheng Guo, Liyong Deng, Badri Vardarajan and Jing He from Columbia University. We also thank Dr. George B Mychaliska and Jeannie Kreutzman from University of Michigan; Dr. Robert Cusick and Sheila Horak from University of Nebraska; Dr. Douglas Potoka, Laurie Luther and Min Shi from University of Pittsburgh; Dr. Gundrun Aspelund and Julia Wynn from Columbia University; Dr. Marc S. Arkovitz from Tel Hashomer Medical Center; Dr. Charles J. Stolar from California Pediatric Surgery Group; Dr. Kenneth S. Azarow from Oregon Health Science Center for enrolling participants. This work was supported by National Institute of Health grant [HD057036] and was supported in part by Columbia University’s Clinical and Translational Science Award (CTSA); grant [UL1 RR024156] from National Center for Advancing Translational Sciences/National Institutes of Health (NCATS-NCRR/NIH) a grant from CHERUBS, a grant from the National Greek Orthodox Ladies Philoptochos Society, Inc. and generous donations from The Wheeler Foundation, Vanech Family Foundation, Larsen Family, Wilke Family and many other families.

### Author Contributions

S.E.C. and T.L. wrote the manuscript, S.E.C. analyzed the data, S.E.C. and P.M. developed the statistical model, W.K.C and Y.S. provided clinical data.

---

**Figure S1.**
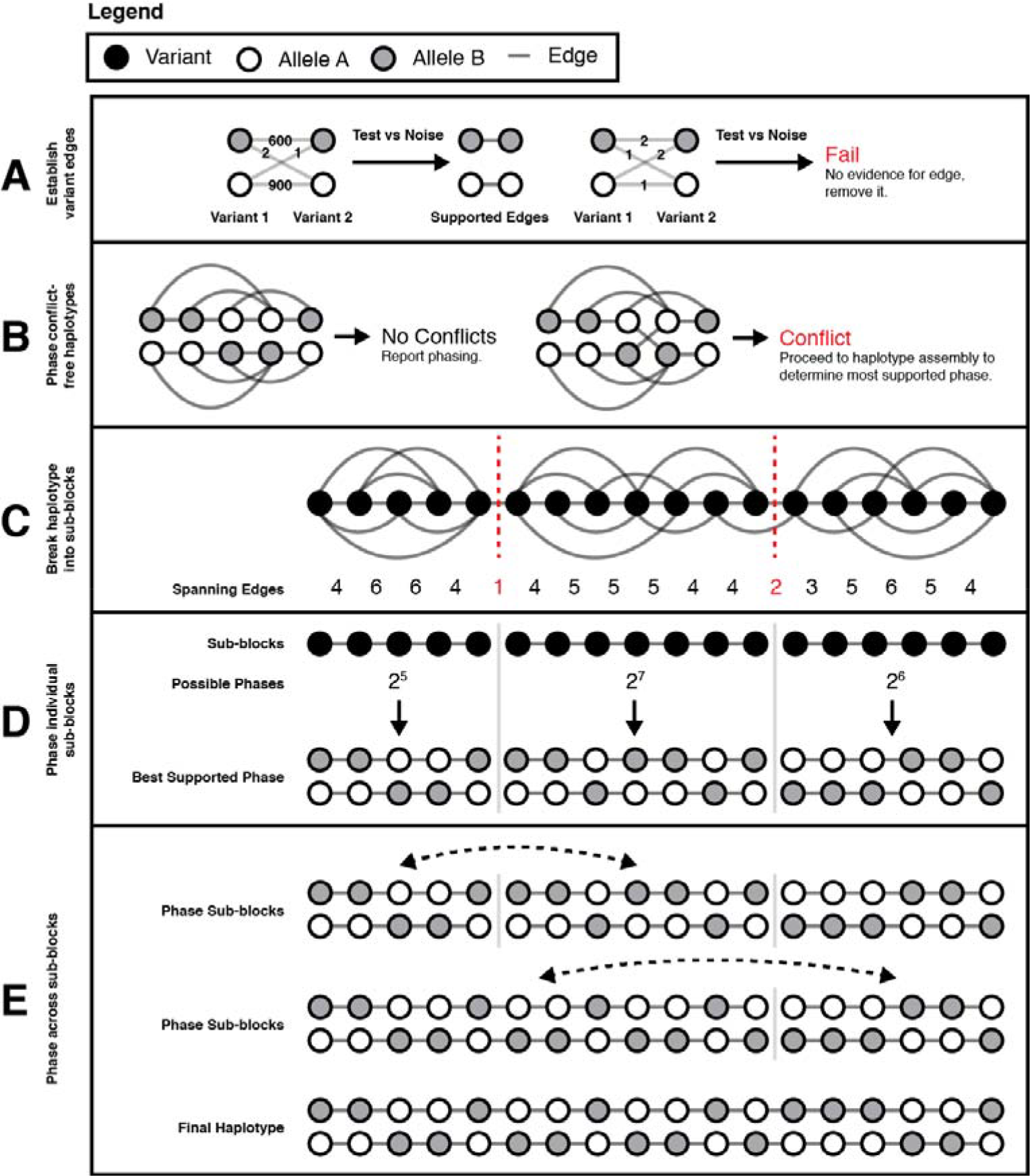
Haplotype assembly in phASER. A) The edges between alleles of each pair of variants that are connected by a read are defined by selecting the phase configuration with the most supporting reads. A binomial test is performed to determine if the number of reads supporting alternative phase configurations can be explained by the amount of sequencing noise in the experiment (Figure S2a). Any variant pairs that fail this test have all edges removed for subsequent haplotype assembly. B) Using the allele edges defined in (A) individual haplotypes are assembled by starting with a single unphased allele and recursively adding all other connected alleles. If after this process two distinct haplotypes arise, where each haplotype contains only a single allele of each variant, the phase is determined to be conflict free and immediately reported. In cases where a single phase is not resolved, haplotype assembly is used to determine the phase most supported by sequencing reads. C) Haplotypes are split into sub-blocks such that they contain no more than a user-defined number of variants per block (shown here for a maximum of 7 per block). Haplotypes are split at positions with the fewest number of edges spanning them. D) Within each sub-block the number of allele edges that support each possible haplotype configuration is tested, and the most supported phase is selected. The number of possible configurations is equal to 2^^^number of variants. E) Once sub-blocks have been internally phased, they are then phased relative to one another, again selecting the configuration with the most support, starting by phasing the two most upstream subblocks with each other, and then subsequently phasing each downstream sub-block until a single haplotype phase is obtained.

**Figure S2.**
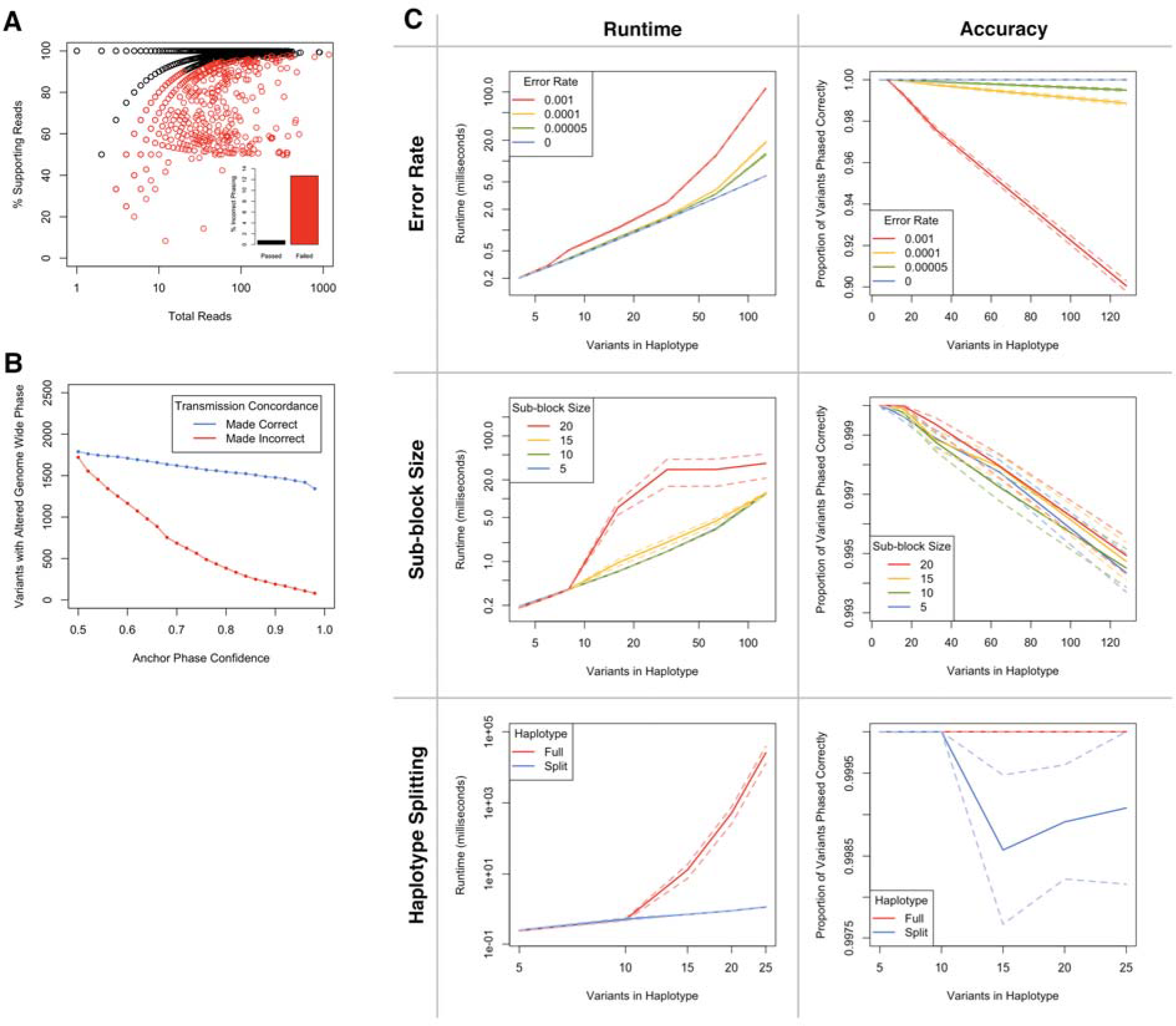
Benchmarking of haplotype assembly and phase anchoring in phASER. A) Percentage of reads supporting the chosen phase as a function of total reads at each variant - variant connection for phASER run using NA12878 RNA ^11^ + WES data ^15^. Each point represents a variant - variant phasing, points in black passed the phase confidence test, while those in red failed (evidence for a conflicting phase configuration, p < 0.01). See “Statistical test for conflicting phasing between two variants” in the methods section for more detail. The inset bar plot shows the percentage of variant pairs phased incorrectly (versus transmission phasing) for variant connections that passed the test (black) and those that failed (red). B) The anchor confidence statistic robustly removes incorrect genome wide phasing as shown by the number of variants with genome wide phase either made correct or made incorrect as a function of anchor phase confidence with phasing by transmission as a ground truth for NA12878 run with 1000 Genomes exome ^15^, Illumina Platinum Genomes WGS (5x) and LCL RNA-seq data ^11^. See “Genome wide phasing using phase anchoring” in the methods section for more detail. C) Benchmarking of the runtime and accuracy of haplotype assembly used in phASER. Observed variant connections within haplotypes of increasing size (defined by number of variants) were simulated 1000 times using defined error rates (probability of edges between the alleles of two variants being incorrect given a known true phase), and with the same distribution of connections across variants within a haplotype as observed in WGS. For reference, the observed error rate of the NA12878 LCL RNA-seq library was calculated as 6.77e-5, so 5e-5 is included as a comparison. Haplotype assembly was then performed using the simulated haplotypes and variants edges with the phASER method, and both runtime and accuracy were reported. This was done while holding sub-block size constant at 10 while varying the error rate, holding error rate constant at 5e-5 and varying sub-block size, and either phasing splitting haplotypes into sub-blocks of 10 variants at most, or without splitting.

**Figure S3.**
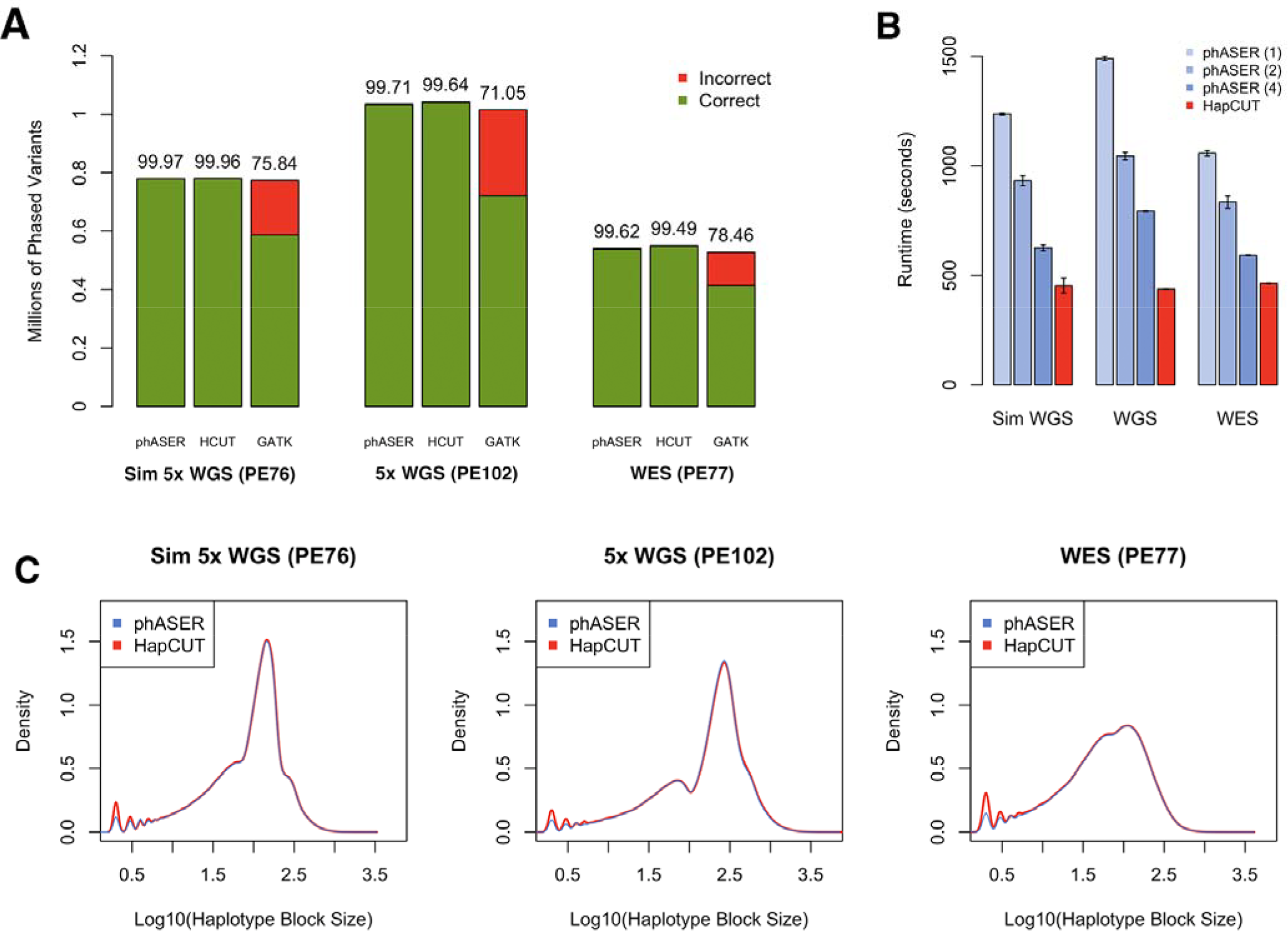
Variant phasing is efficient and accurate using phASER. Comparison of phASER to HapCUT and the GATK Read Backed Phasing tool, using either simulated 5x WGS (NA06986, PE76), experimental 5x WGS (NA12878 Illumina Platinum Genome, PE102), or WES (NA12878 1000 Genomes phase 3 ^15^, PE77). phASER and HapCUT are able to phase a similar numbers of variants, at high accuracy (A), with comparable runtimes (B), and identical haplotype block sizes (C). Runtime for phASER is shown using 1, 2 or 4 threads, since parallelization is available, unlike in HapCUT. phASER’s increased runtime is a result of phASER being designed from the ground up to work with RNA-seq reads, including many features such as additional quality control and allelic expression reporting that are necessary when using this data type, even though they increase overall runtime. Runtime values are means across 4 replicates, and error bars show the standard error of the mean. In our tests, the accuracy of the GATK tool was poor, so it is not shown in subsequent benchmarks. Accuracy was determined by comparing the inferred phase to the known true phase (either the haplotypes used to simulate reads, or the transmission phased NA12878 haplotypes).

**Figure S4.**
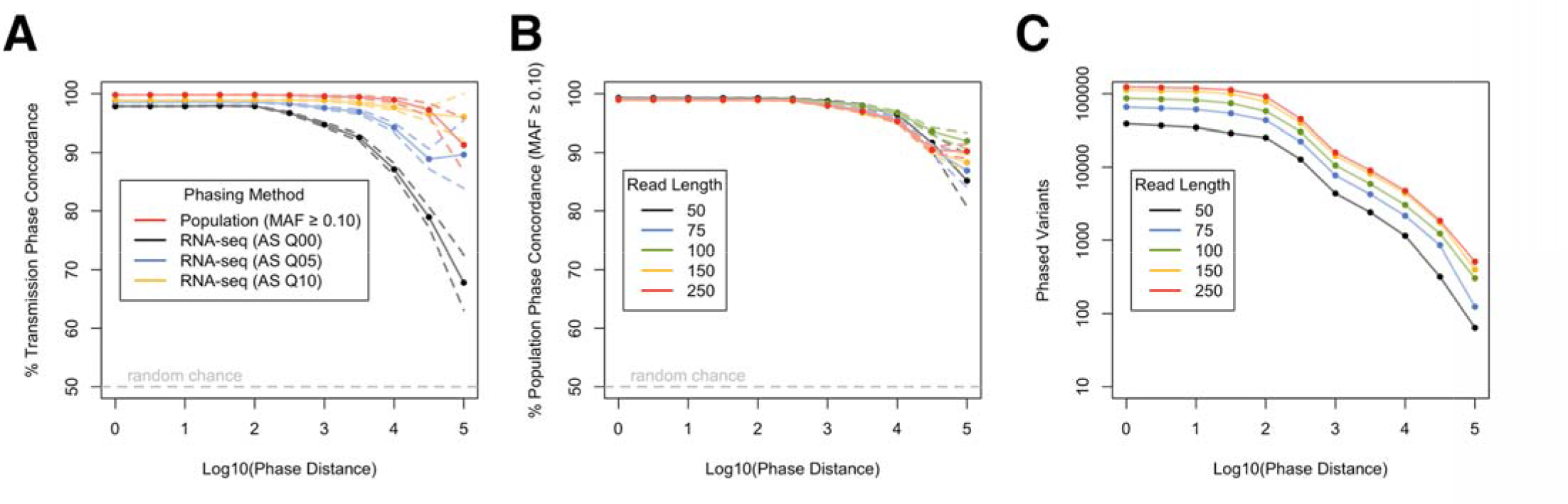
Effects of alignment quality and read length on RNA-seq read backed phasing. A) Comparison of RNA-seq phasing using either no alignment score cutoff (AS Q00), or cutoffs equal to the bottom 5% (AS Q05), and 10% (AS Q10), and population based phasing of common variants (MAF > 0.10) to phasing by transmission. For common variants the accuracy of population phasing is not expected to decrease with distance, so it is shown here as a comparison. Generated using NA12878 LCL RNA-seq data ^11^, 1000 Genomes Phase 3 population phasing ^15^, and Illumina Platinum Genome transmission phasing. B-C) Comparison of RNA-seq phasing to population phasing for variants with MAF > 10% at increasing read lengths (B), and number of variants phased at equal to or greater than increasing genomic distances for increasing read lengths (C), using a GTEx ^12^ long read RNA-seq library (GTEX-WF0N-0001-SM-5S2SE) clipped to various lengths. Solid lines represent the means, and dotted lines the standard error.

**Figure S5.**
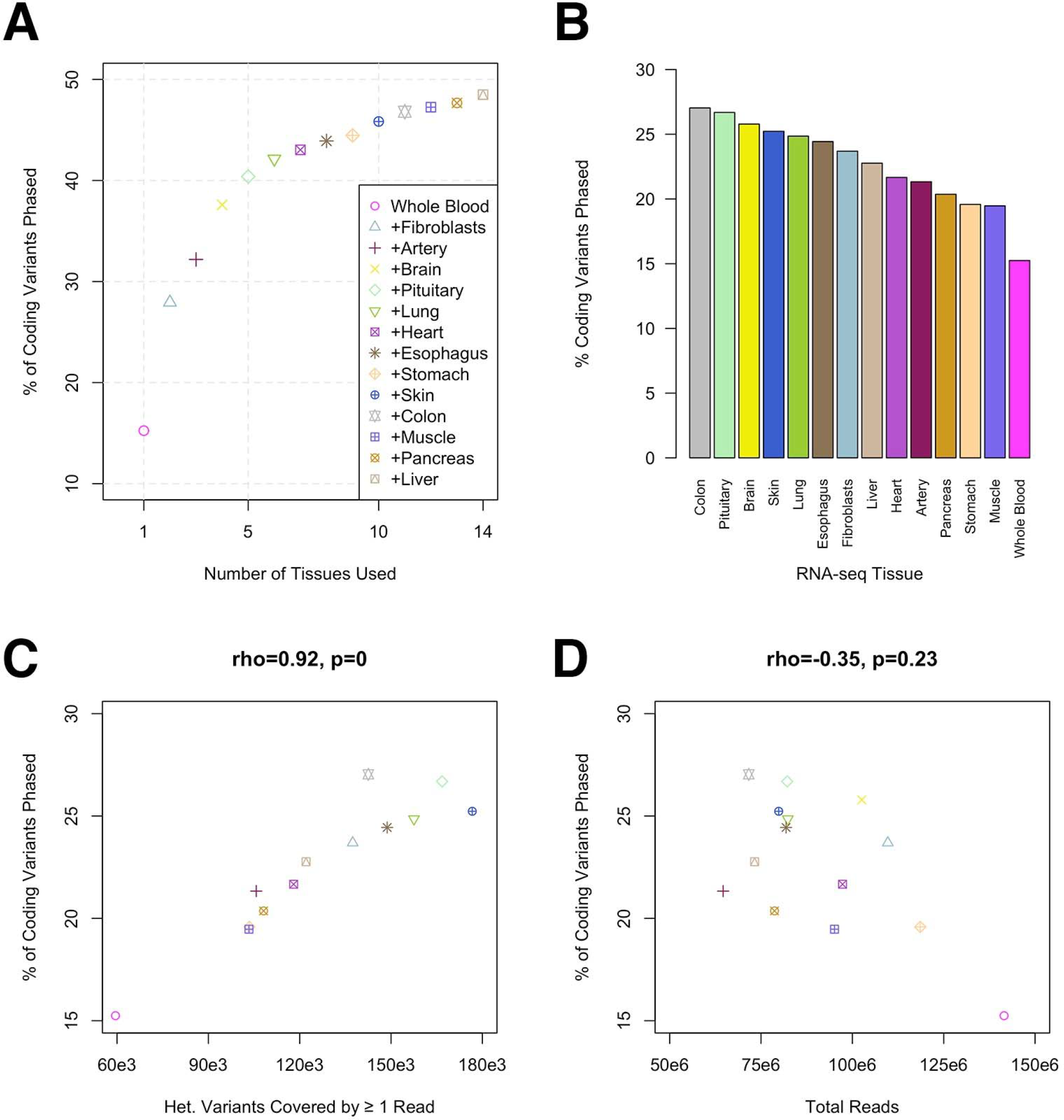
Joint RNA-seq based phasing over multiple tissues greatly improves the number of variants that can be phased in an individual. A) Percentage of coding variants that can be phased beginning with RNA-seq from whole blood, and progressively adding data from up to 14 other distinct tissues from GTEx individual ZAB4 ^12^. B) Percentage of coding variants that can be phased using each tissue from (A) individually. C-D) Percentage of coding variants as a function of the number of heterozygous variants covered by at least one read (C) or total library reads (D) for each tissue from (A) individually.

**Figure S6.**
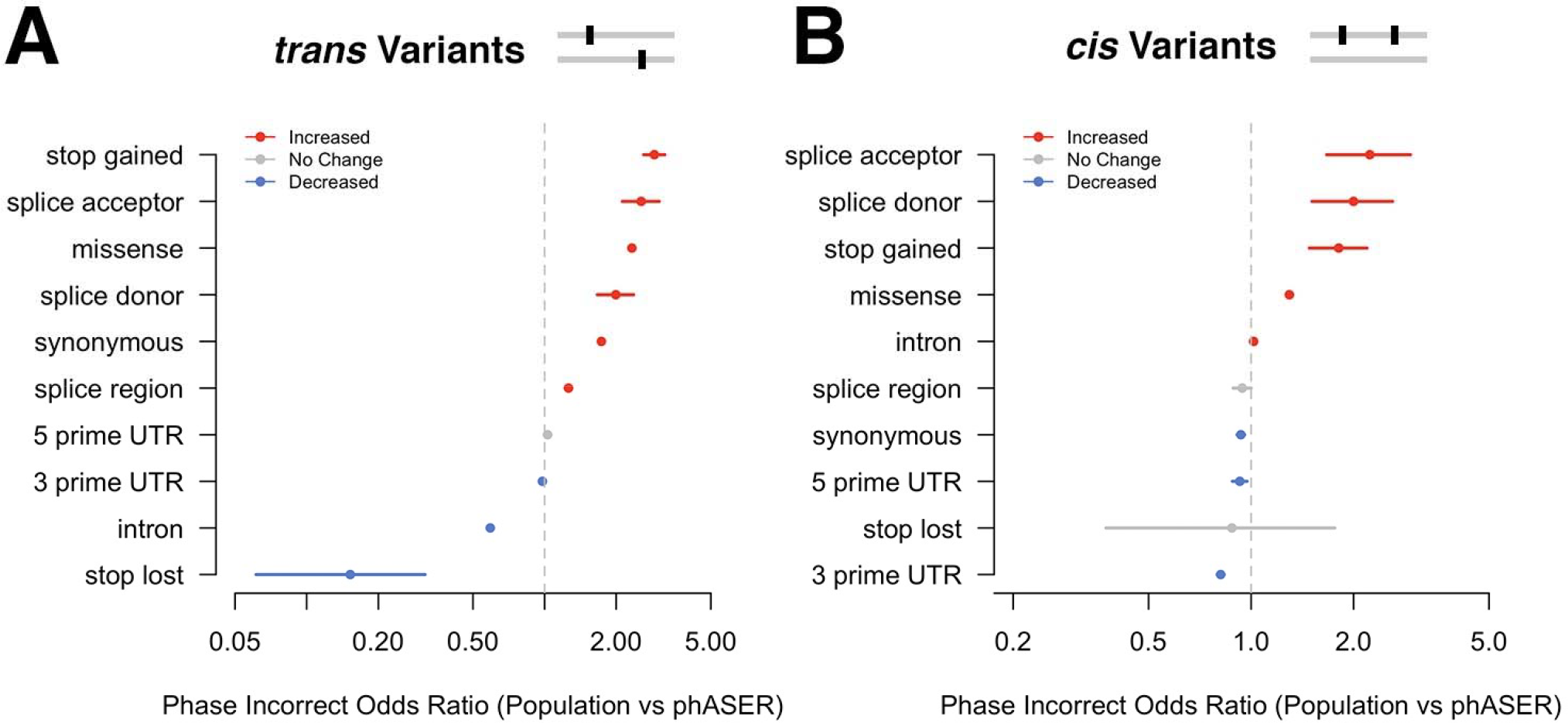
Read backed phasing improves the ability to correctly identify instances of compound heterozygosity at rare variants. Cases of compound heterozygosity were called in 345 individuals using either 1000 Genomes Phase 3 population phasing or exome ^15^ + Geuvadis LCL RNA-seq ^14^ read backed phasing with phASER. The odds ratio for cases involving each variant type being incorrect in population data versus other types in either *trans* (A) or *cis* (B) interactions was calculated using Fisher’s exact test. Variant types with a significantly increased probability of being incorrect when population phasing is used are shown in red, while those with a decrease are shown in blue (p < 0.05). Error bars indicate the 95% confidence interval of the odds ratio.

**Figure S7.**
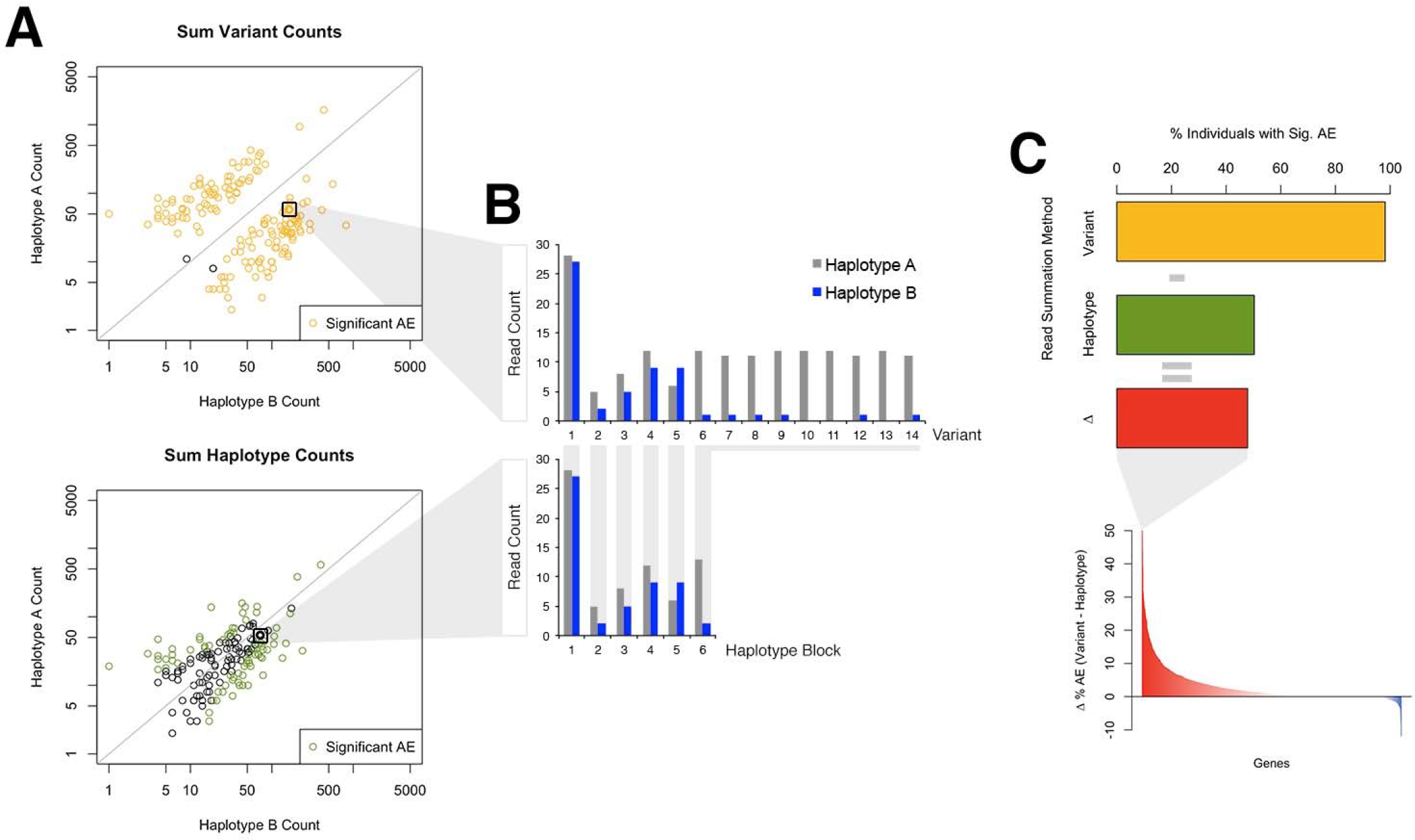
Integrating allelic counts over variants using accurate phasing reduces false positives in allelic expression studies. A) Haplotypic counts for Geuvadis individuals at an example gene (ENSG00000162654 or *GBP4)* calculated by either summing counts from individual variants using 1000 Genomes Phase 3 population phasing ^15^ (top plot, yellow) or with phASER haplotypic counts (bottom plot, green). Each point represents one of the 345 individuals used in the analysis. B) Example illustrating that summing variant counts for the individual highlighted in (A) leads to double counting of variants 7 through 14 in this haplotype (top plot) and is prevented when haplotypic counts are used (bottom plot). C) For this gene the difference (red) in the percentage of individuals showing significant (5% FDR, binomial test) allelic imbalance calculated either by either summing variants (yellow) or using haplotypic counts (green). This value was calculated for all genes with allelic expression data from at least 30 individuals heterozygous for the top Geuvadis cis-eQTL (bottom plot and Figure 2c).

